## Supplementary figures and images for "*Plasmodium* NEK1 coordinates MTOC organisation and kinetochore attachment during rapid mitosis in male gamete formation"

### Fig S1

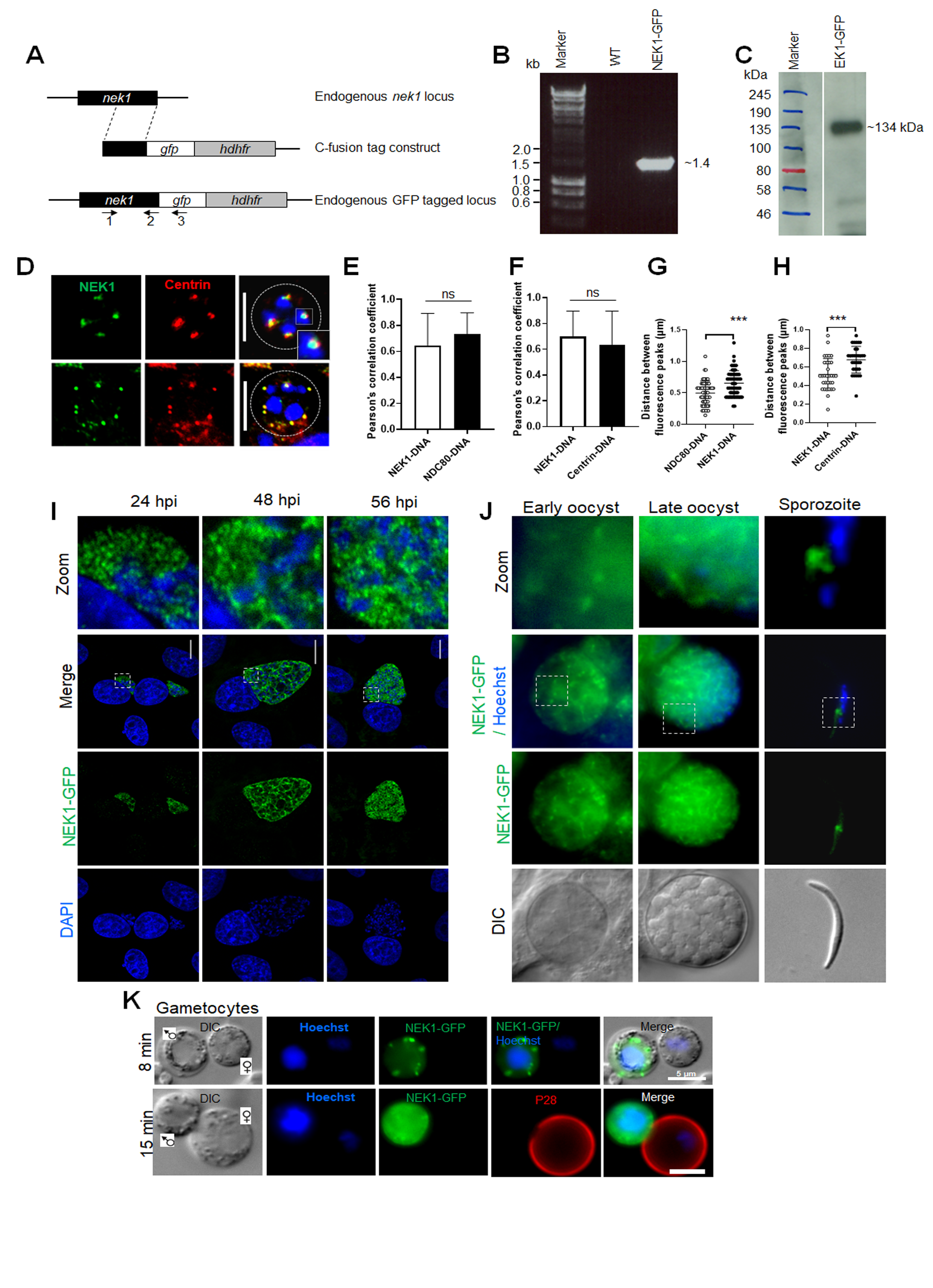

### Fig S2

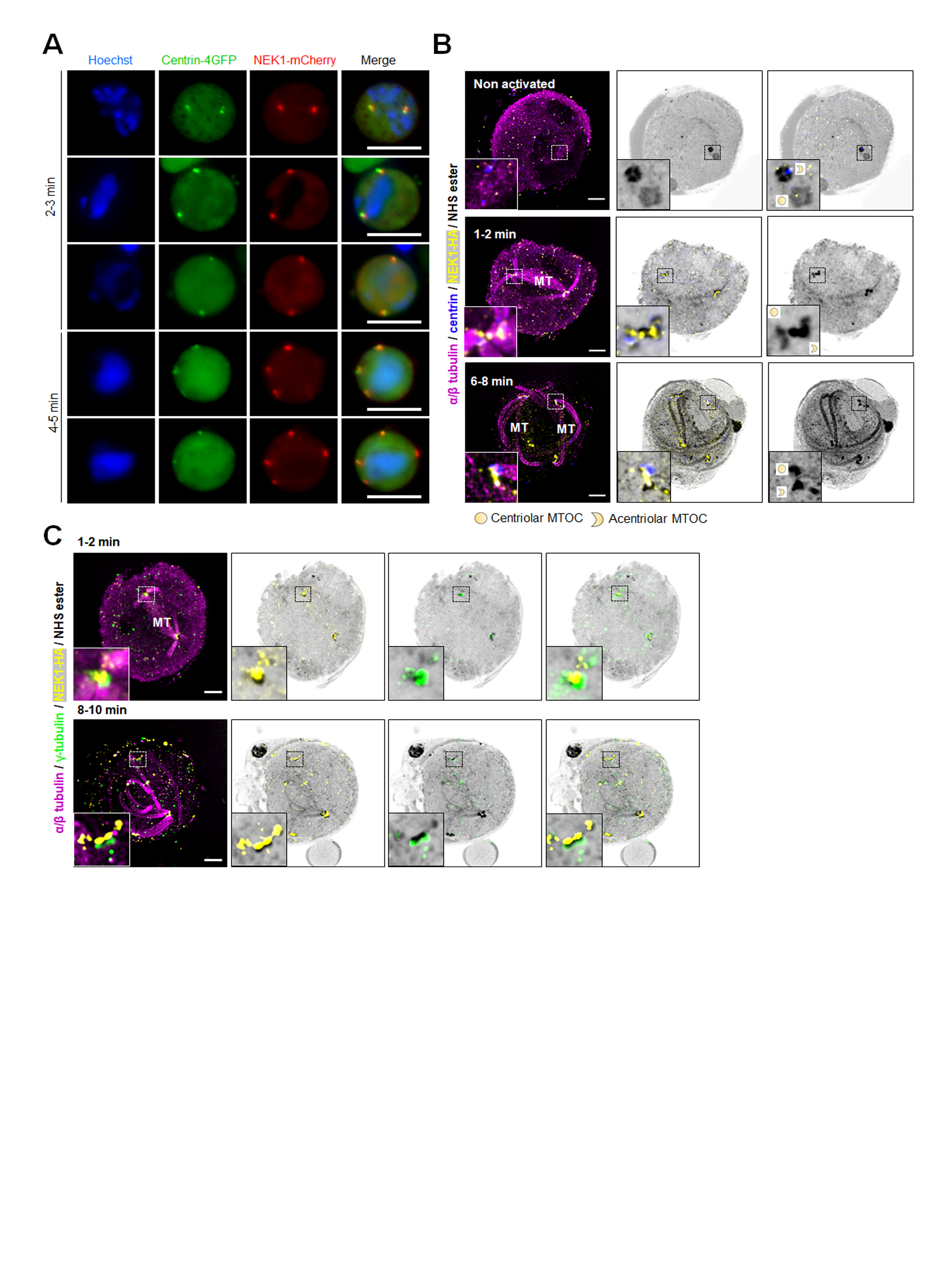

### Fig S3

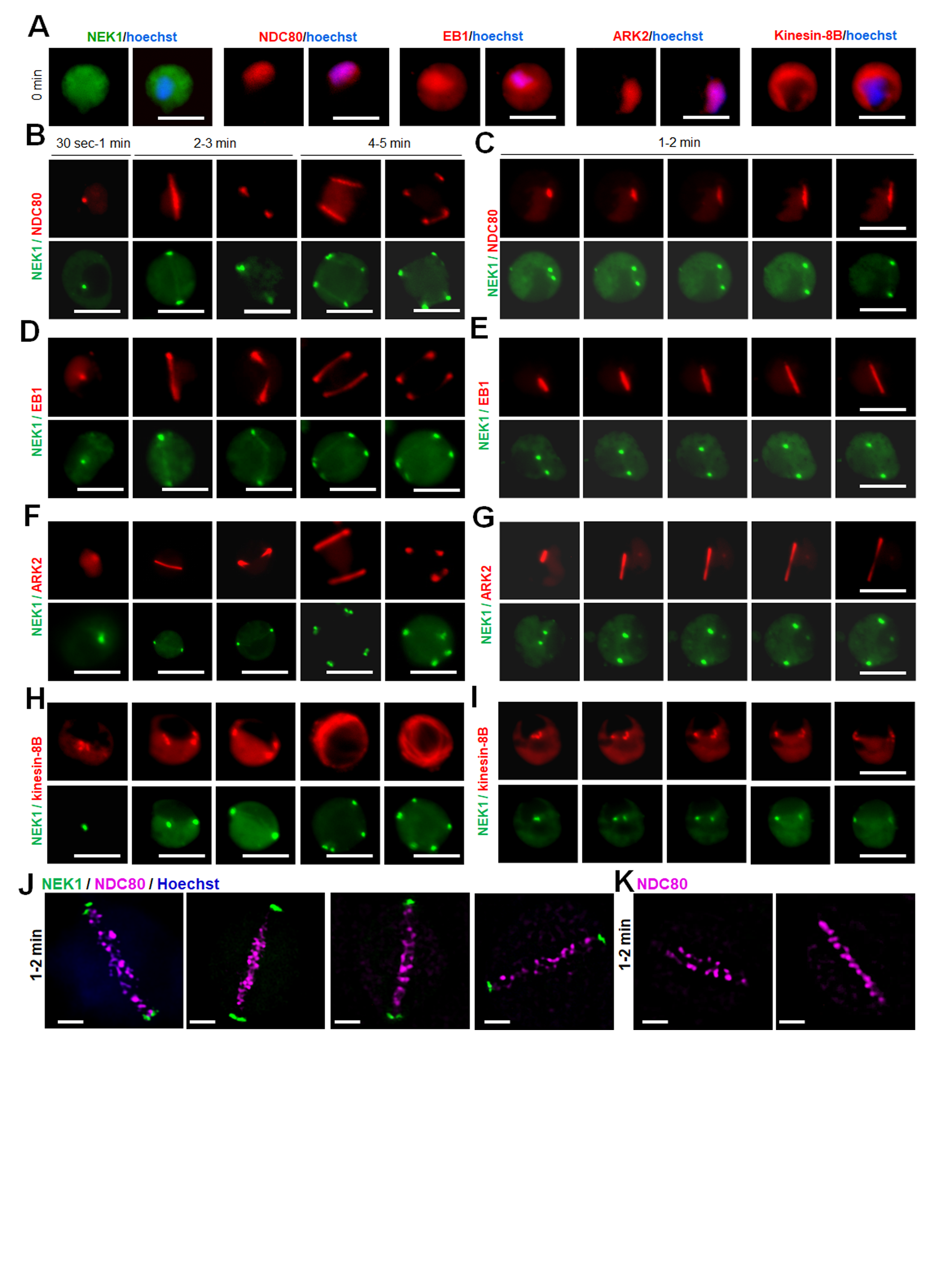

### Fig S4

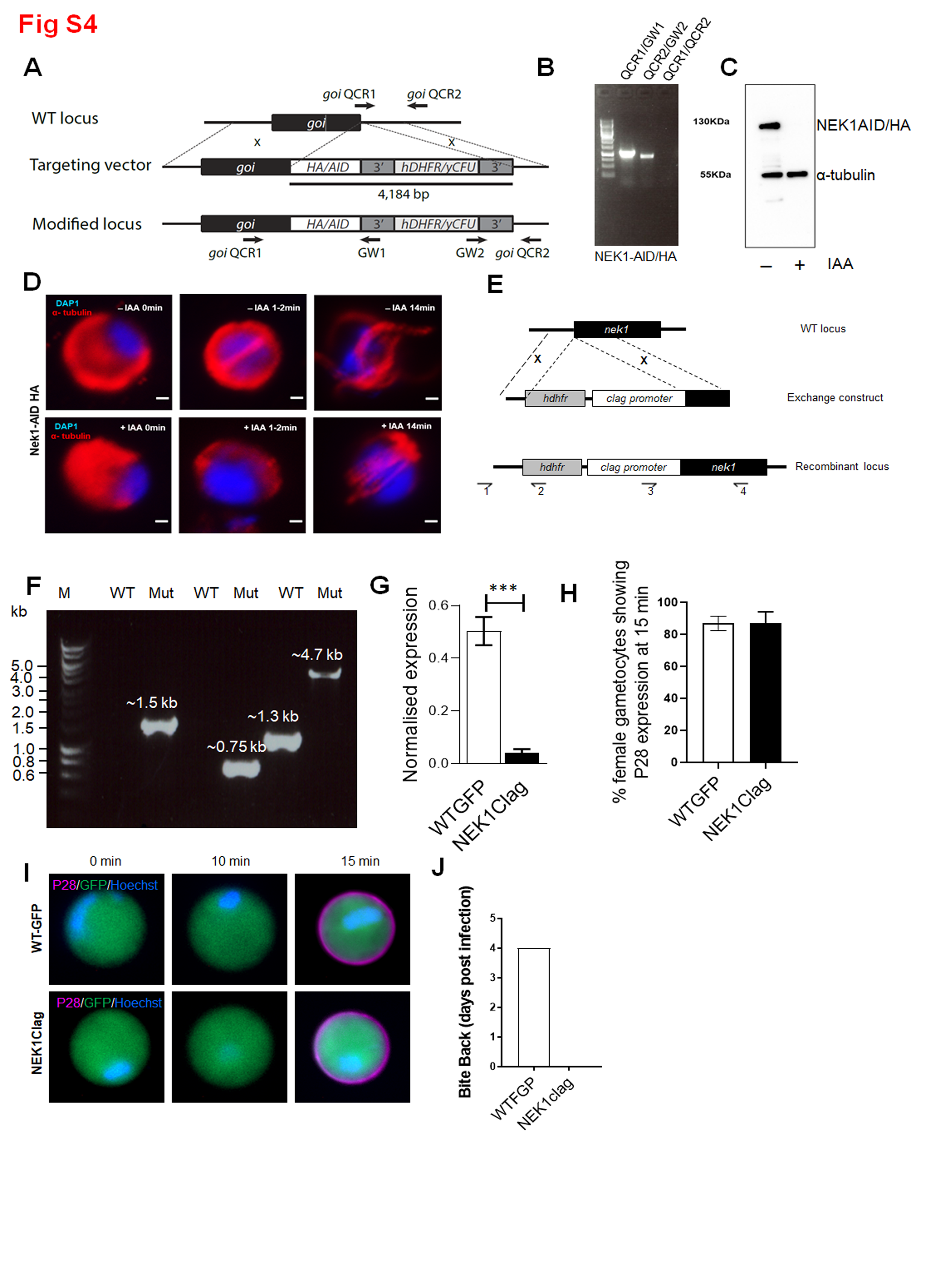

### Fig S5

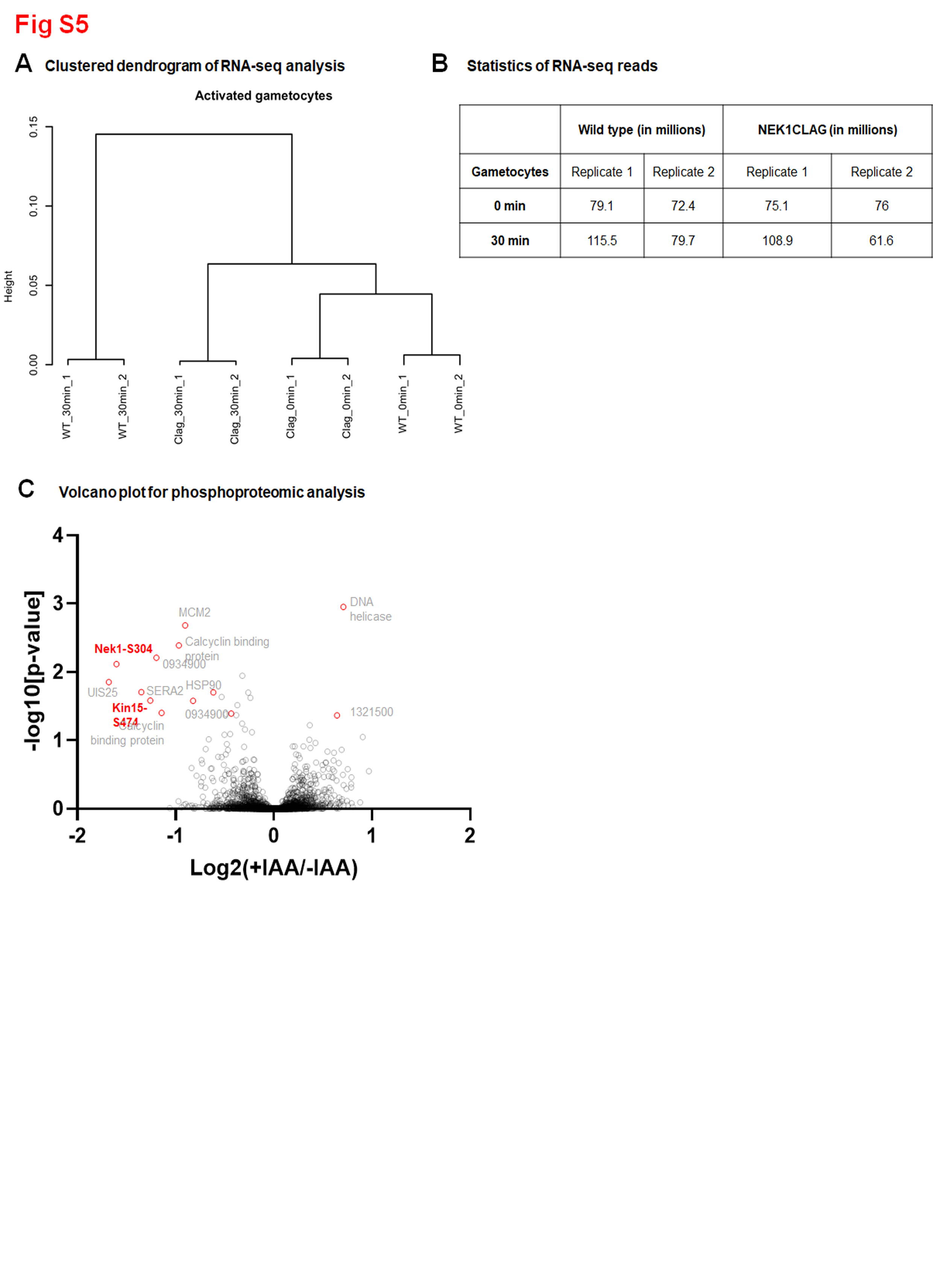

### Fig S6

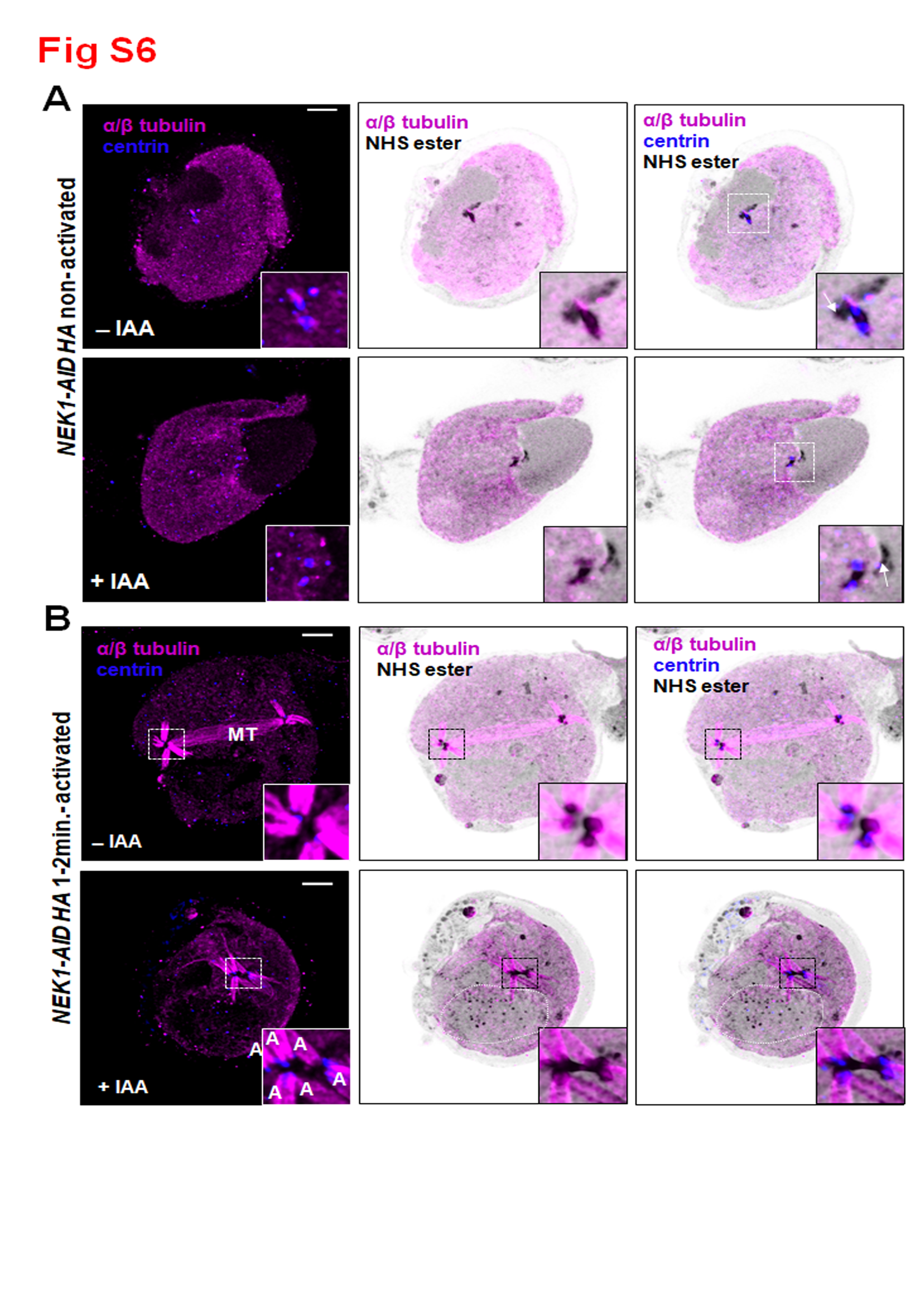

### Fig S7

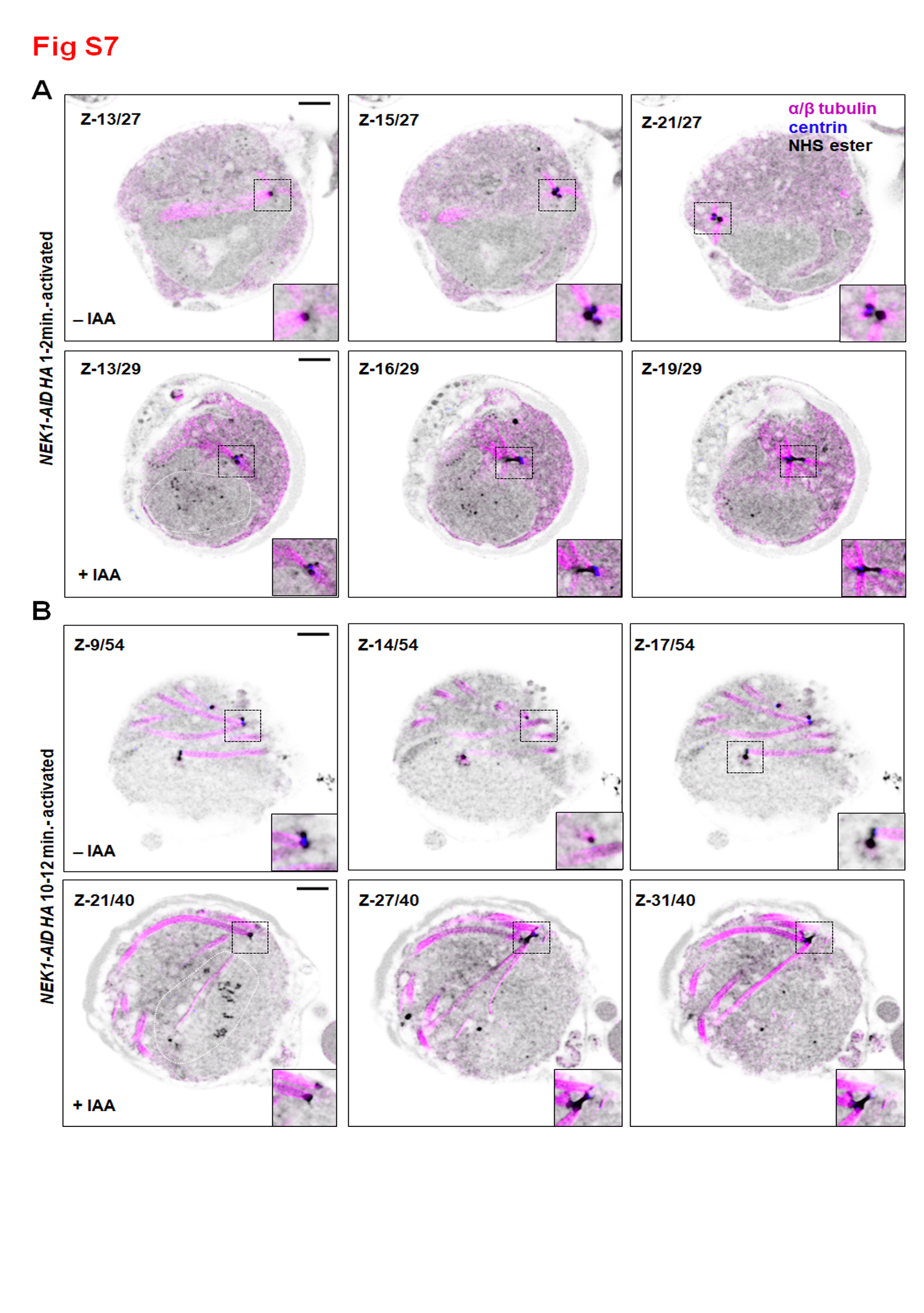
