## Supplementary material for "*Plasmodium* NEK1 coordinates MTOC organisation and kinetochore attachment during rapid mitosis in male gamete formation": Raw images- blot and gels

**Raw images used in the manuscript**

**Agarose gel shown in Fig S1B**

Agarose gel showing the PCR product showing correct integration of GFP at C-terminal of NEK1

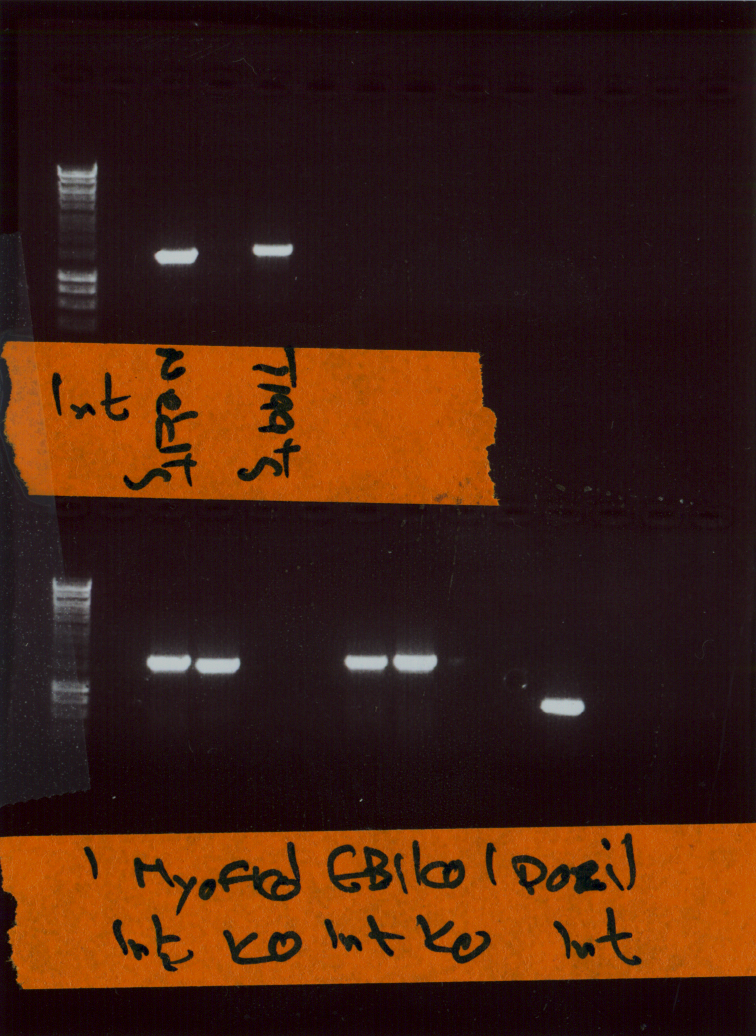

NEK1-GFP

WT

**Raw images-Western blot shown in Fig S1C**

Western blot showing the expression of NEK1-GFP in gametocyte lysate

NEK1-GFP

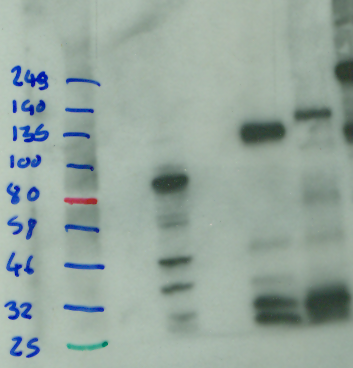
**Raw images-Agarose gel shown in Fig S4B**

Agarose gel showing the PCR product showing correct integration of NEK1-AID

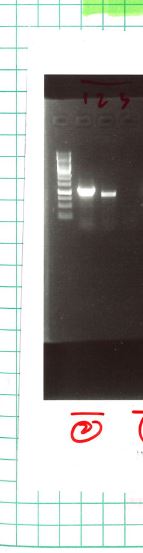

**Raw images-Western blot shown in Fig S4C**

Western blot showing the depletion of NEK1-AID in gametocyte lysate

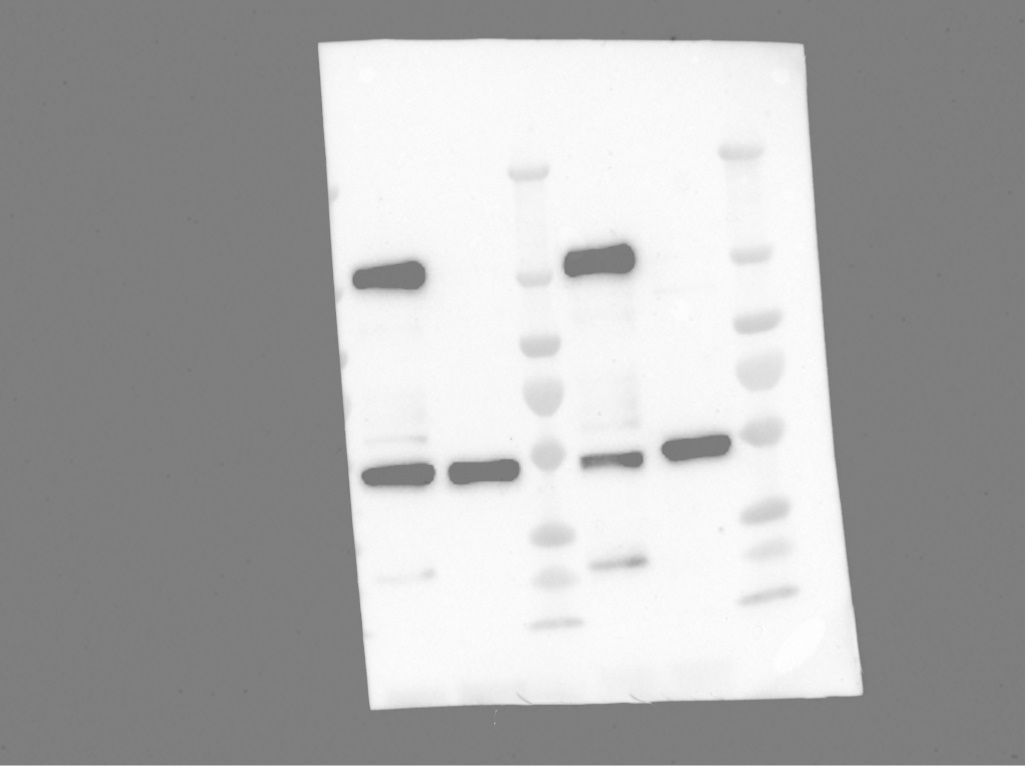

**Raw images-Agarose gel shown in Fig S4F**

Agarose gel displaying the PCR products showing correct integration of NEK1Clag

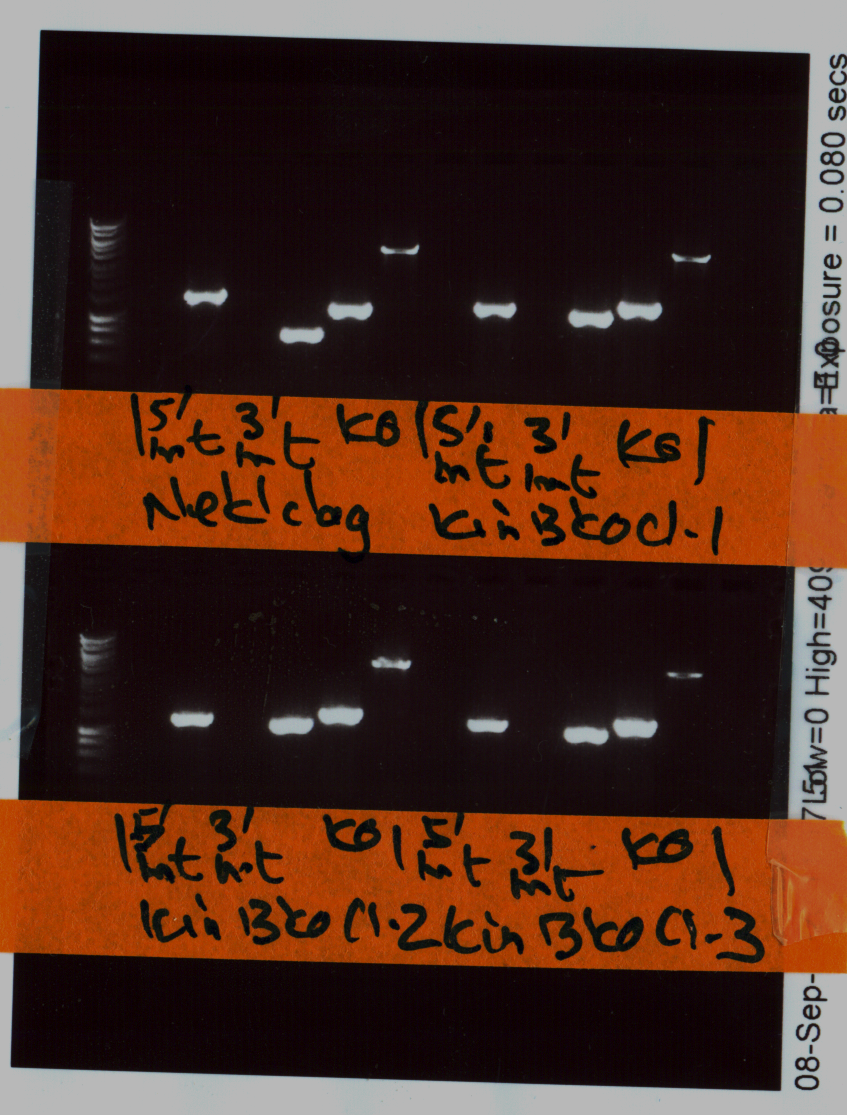
